# Big data approaches to decomposing heterogeneity across the autism spectrum

**DOI:** 10.1101/278788

**Authors:** Michael V. Lombardo, Meng-Chuan Lai, Simon Baron-Cohen

**Affiliations:** Department of Psychology, Center for Applied Neuroscience, University of Cyprus, Nicosia, Cyprus; Autism Research Centre, Department of Psychiatry, University of Cambridge, Cambridge, United Kingdom; Child and Youth Mental Health Collaborative, Centre for Addiction and Mental Health and The Hospital for Sick Children, Department of Psychiatry, University of Toronto, Toronto, Canada; Department of Psychiatry, National Taiwan University Hospital and College of Medicine, Taipei, Taiwan; Cambridgeshire and Peterborough National Health Service Foundation Trust, Cambridge, United Kingdom

## Abstract

Autism is a diagnostic label based on behavior. While the diagnostic criteria attempts to maximize clinical consensus, it also masks a wide degree of heterogeneity between and within individuals at multiple levels of analysis. Understanding this multi-level heterogeneity is of high clinical and translational importance. Here we present organizing principles to frame the work examining multi-level heterogeneity in autism. Theoretical concepts such as ‘spectrum’ or ‘autisms’ reflect non-mutually exclusive explanations regarding continuous/dimensional or categorical/qualitative variation between and within individuals. However, common practices of small sample size studies and case-control models are suboptimal for tackling heterogeneity. Big data is an important ingredient for furthering our understanding heterogeneity in autism. In addition to being ‘feature-rich’, big data should be both ‘broad’ (i.e. large sample size) and ‘deep’ (i.e. multiple levels of data collected on the same individuals). These characteristics help ensure the results from a population are more generalizable and facilitate evaluation of the utility of different models of heterogeneity. A model’s utility can be shown by its ability to explain clinically or mechanistically important phenomena, but also by explaining how variability manifests across different levels of analysis. The directionality for explaining variability across levels can be bottom-up or top-down, and should include the importance of development for characterizing change within individuals. While progress can be made with ‘supervised’ models built upon a priori or theoretically predicted distinctions or dimensions of importance, it will become increasingly important to complement such work with unsupervised data-driven discoveries that leverage unknown and multivariate distinctions within big data. Without a better understanding of how to model heterogeneity between autistic people, progress towards the goal of precision medicine may be limited.

---

Autism occurs in approximately 1-2% of the population^1^ and autistic individuals’ wellbeing is a major public health issue. In economic terms, the lifetime individual cost of autism is estimated at $2.4 (£1.5) million in the USA and UK and annual population costs are around $268 billion in the USA^2, 3^. While interest in and science on autism has been growing rapidly, progress towards translating scientific knowledge into high-impact clinical practice has been less rapid. We are still far from delivering more effective intervention and support, more precise and earlier diagnosis, better understanding and prediction of prognosis and development, and personalization of support and intervention. All of these points are within the scope of stratified psychiatry^4^ and precision medicine^5^. To get to this point, our main contention in this review is that we will first need to grapple with an important issue holding back progress – heterogeneity within the autistic population.

The field is currently grappling with this paramount issue. Some have argued that we are at a crossroads and must acknowledge that the concept of autism as a single entity lacks validity at a biological level^6, 7^ and that autism must be taken apart^8^. This idea relates to what others have discussed regarding autism as an umbrella label referring to many different kinds of ‘autisms’^9^ and regarding how the scientific community should abandon attempts to continue characterizing all of autism under a single theory^10^. Research has begun along these new directions but is highly fractionated because heterogeneity is discussed across multiple levels of analysis, from genetics^11^, neural systems^12-14^, cognition^15^, behavior and development^16-18^, and clinical topics (e.g., response to treatment, outcome^19, 20^). Approaches differ in how heterogeneity is decomposed, from utilizing theoretical *a priori* known stratifiers^21, 22^ or dimensions, to completely data-driven approaches^12, 23-25^. Models for understanding heterogeneity also differ considerably, with some conceptualizing distinctions as categorical/qualitative, continuous/dimensional, and/or where distinctions or similarities may cut across diagnostic boundaries^25-27^. Work can also differ with regards to aims that are specific to understanding heterogeneity within one level of analysis^28, 29^, while others attempt to extend models to explain heterogeneity across levels^30-35^.

The purpose of this paper is not to provide an in-depth review of the literature on these areas. Rather, we see a need to provide organizing principles for framing these diverse areas of research, so that future synthesis and theoretical development about heterogeneity can be facilitated. Specifically, we first discuss the meaning behind the usage of different relevant terminology when discussing heterogeneity in autism. Next, we discuss how heterogeneity arises within the context of the historical change in diagnostic criteria. Third, we provide overarching arguments behind why understanding heterogeneity is critical for furthering progress towards precision medicine. Fourth, we discuss some of the problems with the dominant paradigm still in use in the field – the case-control paradigm. In discussing these issues, we point towards problems with small sample studies and the need for bigger data. This leads into a discussion regarding characteristics of big data that are important for studying heterogeneity in autism. We follow this with organizing principles behind how one attempts to understand multi-level heterogeneity. Finally, we conclude with a discussion about the role of transdiagnostic viewpoints which go beyond understanding heterogeneity just within autism.

## Terminology behind ‘heterogeneity’ and impact on building and evaluating models

The concept of heterogeneity in autism has been around for some time and dates back to the original conceptions of an ‘autistic spectrum’ from Lorna Wing^36^. Since then, we now apply the concept of heterogeneity beyond just clinical, behavioral, and/or cognitive levels. A hallmark of heterogeneity in autism is its multi-level presentation (Fig 1C), applicable from genotype through phenotype^9, 10^ throughout development^16, 37^, and manifesting as important clinical differentiation (e.g., outcome^20^, response to treatment^19^, etc.). Thus, the concept of heterogeneity not only applies to how individuals differ at one level of analysis, but also when and at which levels those differences arise. Theoretically, it is important to consider that multi-level heterogeneity in autism may or may not converge across levels. An important future empirical endeavor will be to sort out how heterogeneous multi-level phenomena may converge or diverge. Pulling such concepts back to familiar language for studying developmental psychopathology, we must understand heterogeneity through the lens of concepts such as equifinality and multifinality^38^. For example, a diversity of different developmental starting points or causal mechanisms in the genome may reach similar endpoints (equifinality) at levels more proximate to clinical outcomes or behavior^39^. However, very similar mechanisms at one level could also result in a diversity of endpoints (multifinality)^40^. Currently the mapping of multi-level heterogeneity in autism is unclear, but it is imperative that we understand these mappings and which are likely to be indicative of useful explanations that place us further down the path towards precision medicine goals.

**Fig 1:**
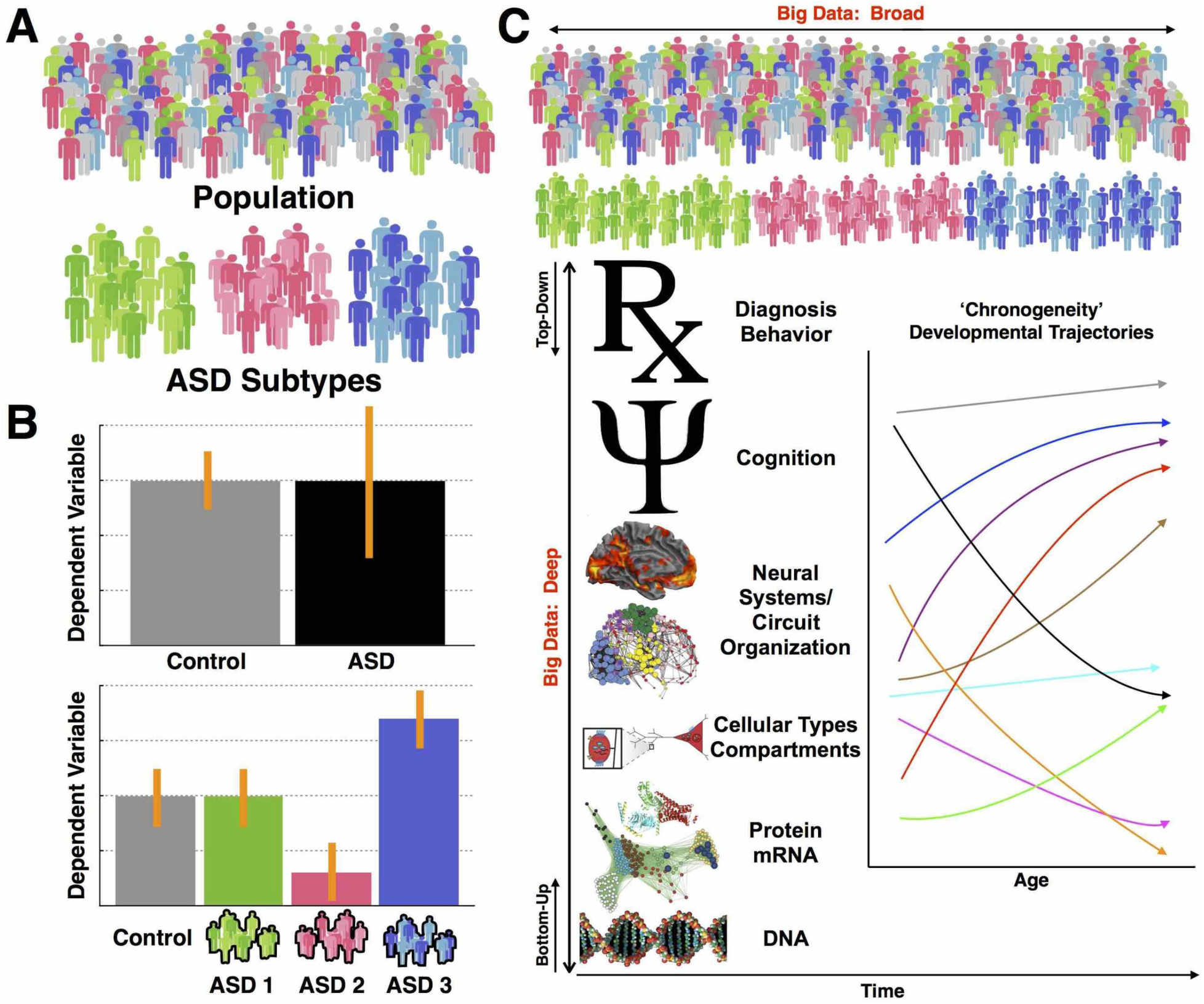
Approaches to decomposing heterogeneity in autism. Panel A shows a population of interest, and autism cases are colored in green, pink, and blue. The different colors are meant to represent different autism subtypes. In panel B we show the impact of ignoring heterogeneity on effect size. With a typical case-control model, we ignore these possible subtype distinctions and compare autism to controls on some dependent variable. In this example scenario there is no clear case-control difference but the autism group shows higher variability (indicated by the larger error bars). An approach towards decomposing heterogeneity might be to construct a stratified model whereby we model the subtype labels instead of one autism label, and then re-examine differences on the hypothetical dependent variable of interest. In this example the autism subtypes show contradictory effects. These effects are masked in the case-control model as the averaging cancels out the interesting different effects across the subgroups. Panel C shows heterogeneity in autism as multi-level phenomena. This panel also visualizes the difference between broad versus deep big data characteristics and labels the top-down versus bottom-up approaches to understanding heterogeneity in this multi-level context. Finally, this panel also shows how development is another important dimension of heterogeneity to consider at each level of analysis (i.e. ‘chronogeneity’). In this example chronogeneity is represented by different trajectories for different types of autism individuals.

There are many ways to talk about how individuals with autism are similar to or different from each other^41^. On the one hand, we can intuitively understand phrases like the *‘spectrum’* as referring to heterogeneity as graded continuous change between individuals. *‘Spectrum’* can also apply to both the clinically diagnosed autism population or the whole population, including those with the ‘broader autism phenotype’^42-45^. The idea of a *spectrum* can be applied as a model for understanding heterogeneity between autistic individuals – a model we would refer to as a ‘*dimensional model*’. Dimensional models can also cut across traditional diagnostic boundaries, with the most prominent example of this being the NIMH Research Domain Criteria (RDoC) model^46^. However, we also use heterogeneity as a way of conceptualizing categorical or qualitative differences between autistic individuals. The term *‘spectrum’* could imply a qualitative, rather than a quantitative difference between individuals. However, terms that pluralize autism as *‘autisms’* may be more applicable here, as the idea of multiple kinds of autisms lends itself to categorical ways of thinking about patients as *‘subgroups’* or *‘subtypes’*. A subtype model for explaining heterogeneity in autism can also be called a ‘*stratified model*’.

Since we have different ways of talking about heterogeneity that have direct impact for how we attempt to build models of within-or between-individual variability, the question will naturally arise as to which way of conceptualizing heterogeneity is best. Are categorical ‘subtype’ models better than continuous ‘dimensional’ models or vice versa? This could be an ill-posed question, since these concepts and models need not be mutually exclusive. First, theoretically we could imagine an important blending between the two types of models for understanding heterogeneity and this can be tested statistically (e.g., factor mixture models^47^). For instance, one could first subtype the autistic population, and then further characterize between-individual variability through continuous models within each subtype. Second, the answer to such a question may differ depending on the aim of the model. For example, a subtype model might be better at predicting treatment responses, whereas a dimensional model might be better at predicting basic biological mechanisms, or vice versa. As we build a literature on understanding heterogeneity in autism, it will be important to be clear about how different models conceptualize heterogeneity, as well as understanding that different models may be important for different types of aims. The statistical aphorism by George Box that *‘all models are wrong, but some are useful’* is applicable here^48^. Models are simplified explanations that typically account only for some portion of variability in a phenomenon. Even if models are quite different in their explanation and predictive power, they can still be quite useful for a variety of different aims. Therefore, pragmatic way of evaluating heterogeneity models is important for moving forward, since it is unlikely that we’ll converge on single explanations (models) that can explain the wide array of multi-level heterogeneity in autism.

## Heterogeneity, evolution of the diagnostic concept

The evolution of the nosology and diagnostic concept of autism inevitably changes the definition of autism – who counts as being on ‘on the spectrum’ and who gets a clinical diagnosis^49^. This evolution also inevitably contributes to the discussion about heterogeneity in autism. When ‘autism’ was first defined as ‘autistic disturbances of affective contact’, the core features were considered to be ‘extreme self-isolation’ and ‘obsessive insistence on the preservation of sameness’^50, 51^. At the cognitive level, language impairments or peculiarities were seen as secondary to ‘basic disturbances in human relatedness’^51^. Moreover, both Leo Kanner and Hans Asperger recognized good cognitive potential in their child patients^50, 52^ and therefore autism was not necessarily tied to intellectual disability. However, at the next stage of nosological evolution, language and cognitive impairments began to be considered ‘core’^53^ and this conceptualization directly impacted the first operationalization of autism in the DSM-III^54^, in which language deficits were core to diagnosis. Individuals identified as having autism in the 1970s and 1980s were therefore mostly those with marked difficulties in verbal communication, and many were considered to have intellectual disability. In the 1980s, Lorna Wing and colleagues not only introduced the work of Hans Asperger into the English speaking world^55^ but also conducted epidemiological studies that demonstrated the heterogeneity in social, language, motor, and cognitive abilities in the autistic and developmentally delayed population^56, 57^. Wing’s ideas of the ‘triad of social and language impairment’, the lack of clear division between Kanner’s autism and less extreme forms, and the shift of core social impairment from ‘extreme autistic aloneness’ to ‘deficits in the use and understanding of unwritten rules of social behavior’, clearly broadened what autism encompassed. All these ideas were subsequently adopted into versions of diagnostic systems including DSM-III-R, DSM-IV and ICD-10. Phenotypic heterogeneity therefore increased, allowing an autistic individual to be verbal or minimally verbal, ‘active but odd’, ‘passive’, ‘aloof’ or ‘loners’^58^, and with various combinations of repetitive and stereotyped behaviors. The DSM-5’s exclusion of language impairments from, and inclusion of sensory idiosyncratic responses into core symptoms, reflects how the concept of autism nowadays is much broader than how it had initially been conceptualized. With the changing and broadening diagnostic concept comes increased heterogeneity, inevitably at the behavioral phenotypic level, and possibly also at other levels of analysis.

This history behind the evolving diagnostic concept is an important, yet often not fully acknowledged caveat for interpreting research on autism. Research spanning several decades may have been isolating phenomena in altogether different types of individuals than does more modern research. Since the spectrum of diagnosed individuals is wider today than in the past, interpretations behind lack of replication or inconsistencies across studies should take this into account, rather than assuming the population under study has not changed over time. As the diagnostic concept continues to change we must be mindful of this issue when interpreting how current research matches up to work that may be several decades old.

## Shifting from the ‘one-size-fits-all’ paradigm towards understanding heterogeneity

Perhaps the most prominent justification behind why understanding heterogeneity is important is because individuals with autism widely differ in response to treatment. While most treatment approaches are early intensive behavioral intervention (EIBI) and naturalistic developmental behavioral intervention (NDRI), the existing literature suggests that they have variable levels of effectiveness and in some cases may not significantly affect core autism features such as social-communication difficulties^59-63^. Currently there are also no medical treatments that significantly affect the core characteristics of autism^64, 65^. Rather than advocating a ‘*one-size-fits-all*’ approach to treatment, most recent best practice recommendations specifically highlight the critical need for future research to identify factors that explain heterogeneity in response to treatment, in order to better individualize treatment approaches and to better target changes in core symptomatology^59, 63^.

Heterogeneity also limits basic scientific progress towards understanding autism. To understand why, it is important to first make salient the problems with the dominant paradigm, which is ill-equipped to clarify heterogeneity - the case-control paradigm. The case-control paradigm exemplifies the ‘*one-size-fits-all*’ approach, since all cases are treated identically due to the same diagnostic label. Studies that attempt to identify ‘biomarkers’ via case-control designs have implicitly conceptualized the notion that if a strong biomarker did exist, that it would completely differentiate cases from all controls. We have yet to isolate any biomarkers for autism that can reliably and consistently reach this high bar^7, 66^. One reason why case-control research has fallen short on identifying high impact biomarkers could be that we are looking at the wrong features. However, an alternative explanation is that high-impact biomarkers are likely exclusive to specific subsets of autistic individuals. That is, a high-impact biomarker may be informative for one subtype of autism, but not others (Fig 1B). In order to identify such stratification or dimensional biomarkers^67^, one will have to change the approach from the case-control model to a stratified and/or dimensional model. This is not to say that case-control studies are not useful. Isolation of consistent and reliable case-control differences are useful for identifying on-average differences, but typically with substantial degree of overlap in the distributions. However, because we are in the search for biomarkers that could help us move towards precision medicine, we will need to pivot our approach away from case-control studies as the dominant paradigm and towards stratified models that could yield much higher impact larger effects.

As an illustrative example, we take our own recent work on mentalizing ability in adults with autism. From a case-control perspective, adults with autism perform on-average lower on the ‘Reading the Mind in the Eyes’ Test (RMET) compared to matched typically-developing controls^68^. However, taking a stratified approach, we find that the autistic adult population can be reliably split into subtypes that either are completely unimpaired on the RMET, versus those who are highly impaired^24^ (Fig 2). Thus, in this example, while replicable on-average case-control effects appear, a stratified approach that takes into account heterogeneity can isolate higher impact and more precise considerations about mentalizing as measured by the RMET in the adult autistic population.

**Fig 2:**
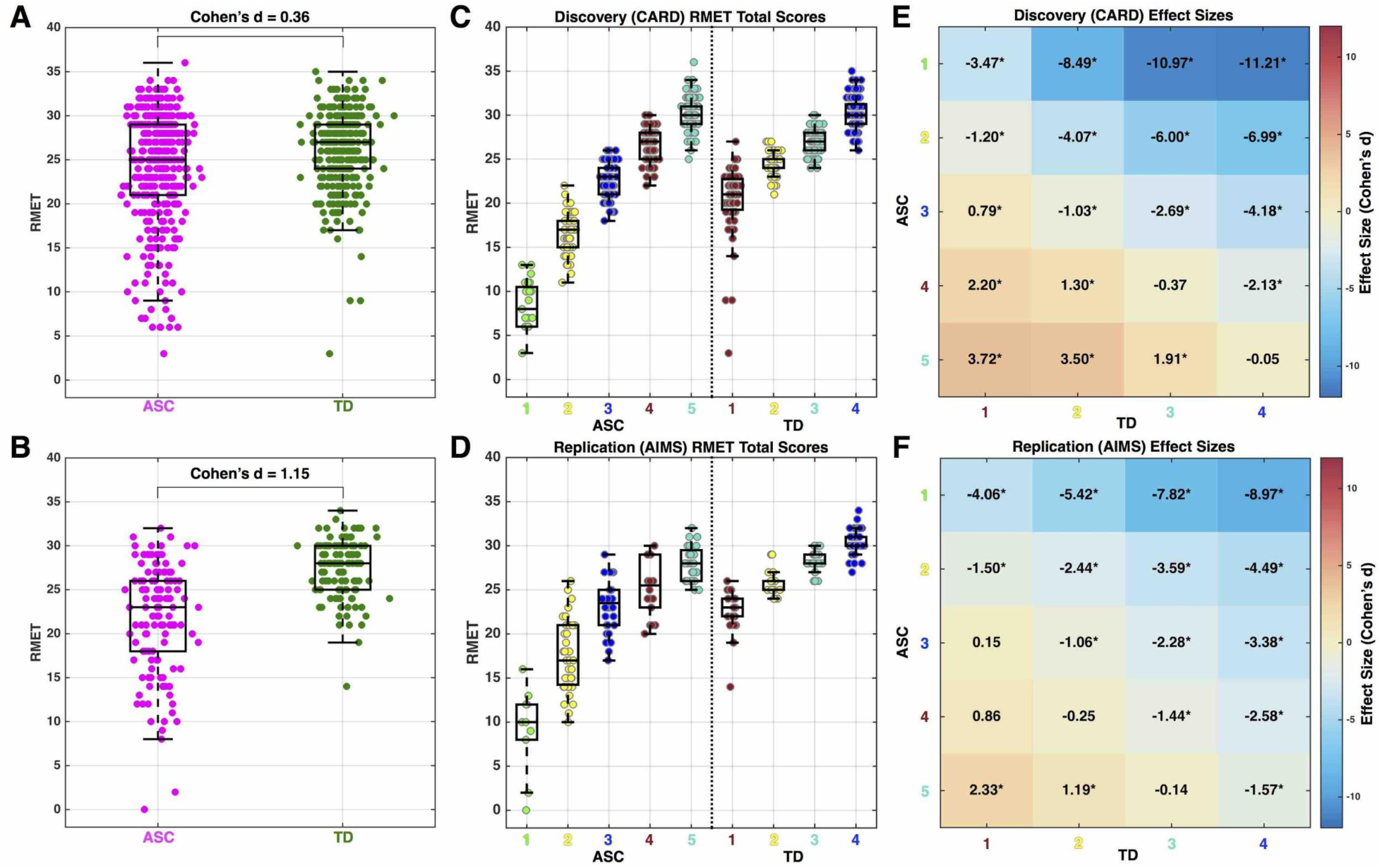
Case-control vs stratified model example with adult autism and mentalizing ability. This figure reports data from Lombardo et al., (2016)^24^ on two independent datasets of adults with autism and performance on an advanced mentalizing test, the Reading the Mind in the Eyes Test (RMET). Panels A (Discovery) and B (Replication) show case-control differentiation and the standardized effect size for each dataset. Panels C-F shows RMET scores and standardized effect sizes from the same two datasets after unsupervised data-driven stratification into 5 distinct autism subgroups and 4 distinct TD subgroups. autism subgroups 1-2 are highly impaired on the RMET, while autism subgroups 3-5 are completely overlapping in RMET scores with the TD population.

## Imprecise effect size estimates and lack of power in small sample size studies

Compounding the problem of utilizing ‘*one-size-fits-all*’ models like the case-control paradigm is the issue of small sample size studies. Over the last several decades, it has been common practice to conduct and publish small sample size studies. Small sample studies can be problematic from the statistical viewpoint that statistical power is low for all but the largest effects. Small sample size also means that estimated sample statistics vary considerably relative to their population parameters in small samples due to more pronounced sampling variability. In Fig 3, we show simulations that illustrate the issues of low power and imprecise estimates of effect size so that they are clear and salient to readers. A common case-control study with n=20 per group results in an effect size that varies considerably relative to the true population effect. This variability in estimated effect size at small samples is consistent irrespective of what the true population effect is. Only with very large sample sizes (e.g., n>1000) can we see that sample effect size hones in with some precision on the true population effect size. The histograms shaded in red in Fig 3 also show the limited statistical power one has at smaller effect sizes and small sample size.

**Figure 3:**
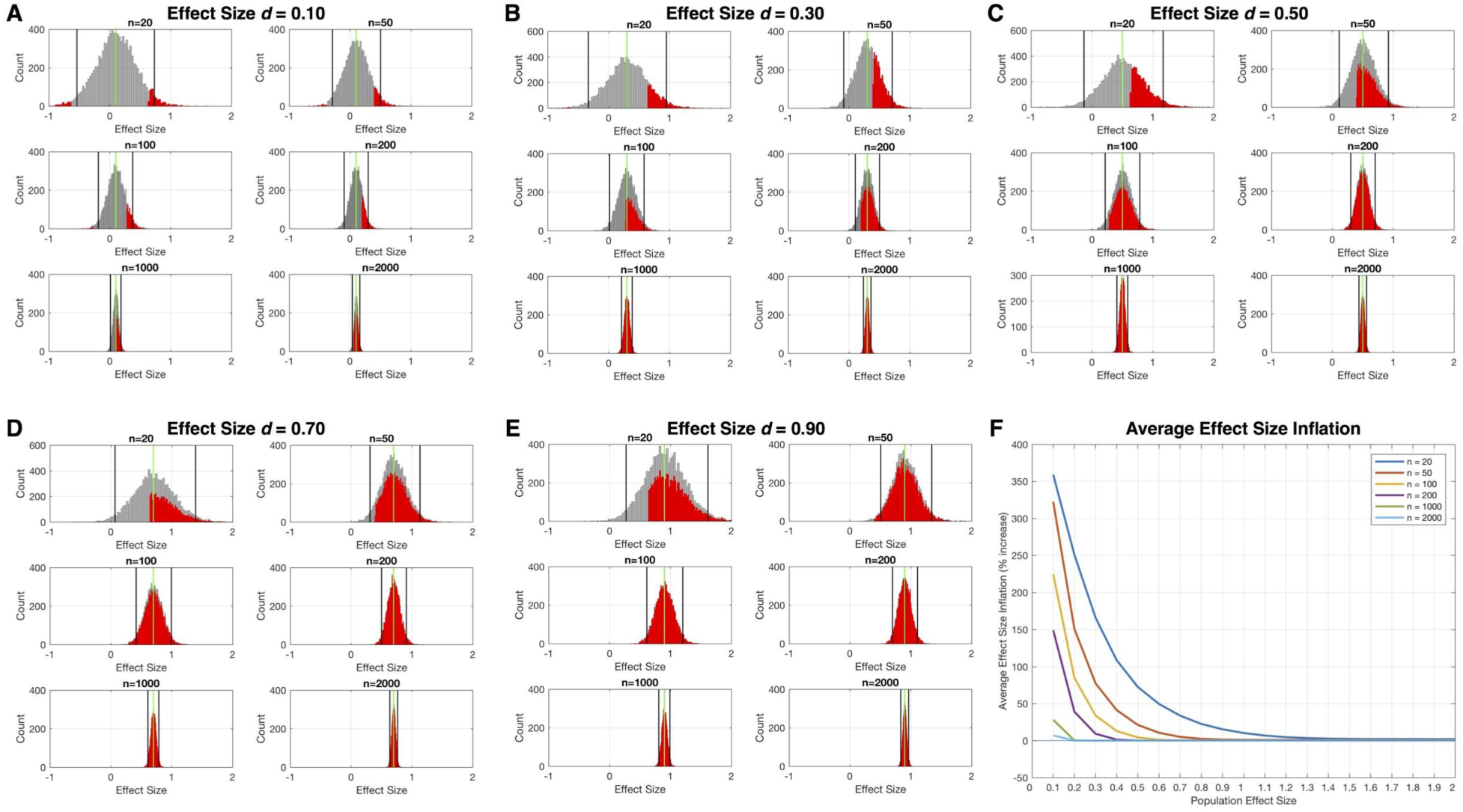
Simulation of sample effect size estimates at different sample sizes and across a range of true population effects for a hypothetical case-control study. In this simulation we set the population effect size to a range of different values, from very small (e.g., d=0.1) to very large (e.g., d>1.0). We then simulated data from two populations (cases and controls), each with n=10,000,000, that had a case-control difference at these population effect sizes. Next, we simulated 10,000 experiments where we randomly sampled from these populations different sample sizes (n=20, n=50, n=100, n=200, n=1000, n=2000) and computed the sample effect size estimate (standardized effect size, Cohen’s d) for the case-control difference. These histograms (grey) show how variable the sample effect size estimates are (black lines show 95% confidence intervals) relative to the true population effect size (green line). Visually, it is quite apparent how small sample sizes (e.g., n=20) have wildly varying sample effect size estimates and that this variability is consistent irrespective of what the true population effect size is. Overlaid on each grey histogram are red histograms that show the distribution of sample effect size estimates where the hypothesis test (e.g., independent samples t-test) passes statistical significance at p<0.05. The rightward shift in this red distribution relative to the true population effect size (green line) illustrates the phenomenon of effect size inflation. The problem is much more pronounced at small sample sizes and when true population effects are smaller. We then computed what is the average effect size inflation for this red distribution and plotted this average effect size inflation as a percentage increase relative to the true population effect in panel F. This plot directly quantifies the degree of effect size inflation across a range of true population effects and across a range of sample sizes. The code for implementing and reproducing these simulations is available at https://github.com/mvlombardo/effectsizesim.

## Effect size inflation in small sample studies

Our simulations also make salient another common characteristic of small sample size studies - the possibility for vast effect size inflation when statistically significant effects are identified^69^. Inflated effects occur because effect sizes that are deemed statistically significant in small studies benefit from noise in the direction of the effect. Such inflated effects present an over-optimistic view on the identified effects and are prone to the winner’s curse^69^. Inflated effects look attractive and may be easier to publish due to their apparent indication of large effects. However, in subsequent replication attempts, investigators likely will fail to identify effects as large as the original small sample study because the effect size in the original study was inflated by some degree. We can clearly see effect size inflation and its interaction with true population effect size in Fig 3. At very small true population effect sizes, sample effect size estimates that are deemed statistically significant (i.e. red histograms in Fig 3A-E) are wildly inflated, and this problem is most pronounced for small sample size studies. For example, tiny population effect sizes of 0.1 standard deviations of difference show on-average greater than 300 to 350% effect size inflation when a study observes a statistically significant effect at p<0.05 with an n=50 or n=20, respectively (Fig 3F). If the true population effect size is much larger (e.g., d>0.5), inflation in effect size is attenuated, and at relatively large sample sizes (n>100 per group), there is very little effect size inflation on-average for such effects. Of course, these simulations here are a simplistic example of a study with only one statistical comparison. The reality is that typically studies make many multiple comparisons and sometimes on a massive scale (e.g., neuroimaging, genetics). In these situations, inflated effect sizes become an even larger problem^70^.

Why is such a characteristic important in discussions on case-control paradigms versus paradigms that acknowledge heterogeneity? The pervasiveness of small sample sizes and effect size inflation in case-control studies tend to give over-optimistic views on the utility of case-control studies. Over the course of time, replication attempts typically decrease the enthusiasm for many such effects, because the reality is likely that most case-control effect sizes are much smaller than published small sample size studies would suggest. By portraying initial novel case-control studies as showing large effects, we may be less inclined to ask the question of whether heterogeneity is involved. Furthermore, very small case-control effects may be due to complicated heterogeneity in the autism population that hides potentially very large effects restricted to specific subtypes. By focusing on heterogeneity, we are likely to better identify true population effects of much larger magnitude. Assuming that such work identifies true large effects in relatively large samples, the issue of effect size inflation may be much less of an issue (as the simulations here demonstrate). However, any models where statistical power is low can show inflated effect sizes. Therefore, models that try to explain heterogeneity can be prone to effect size inflation as well, hence the need for very large samples and high statistical power in stratified or dimensional models.

## Sampling bias across strata nested in the autism population

Small sample size case-control studies that do not acknowledge heterogeneity in the autism population are also particularly problematic because increased sampling variability has substantial biasing impact in enriching specific strata of the population over others. Ideally, to get a generalizable sample of the population in a case-control paradigm, one hopes that if there are unknown strata nested in the population, the sample prevalence of each strata reflects the true prevalence of that strata in the population. If such a criterion is not achieved, it means that samples can be biased by the enrichment of certain strata of the population over others. If enrichment of different strata of the population are present across multiple studies, they may paint a confusing and potentially contradictory picture of the phenomenon. A primary example of this is the systematic over-enrichment of males over females in most case-control studies, particularly intervention and biological studies^71-73^, which may lead to male-biased inferences about autism^74^. Another simulation shown in Fig 4 illustrates that small samples are much more prone to this bias due to enrichment of specific strata over others. In this simulation, there are 5 subtypes in the autism population, and each have different effects relative to the control population. Therefore, enrichment of different subtypes can have dramatic effects on the results of the study. Our simulation had equal population prevalence for each subtype (i.e. 20% in all autism population), which meant that from study to study, the specific strata that may be enriched is random. Obviously, in the likely scenario where population prevalences are asymmetrical across subtypes, the enrichment of specific strata could favor those subtypes with higher population prevalence.

**Figure 4:**
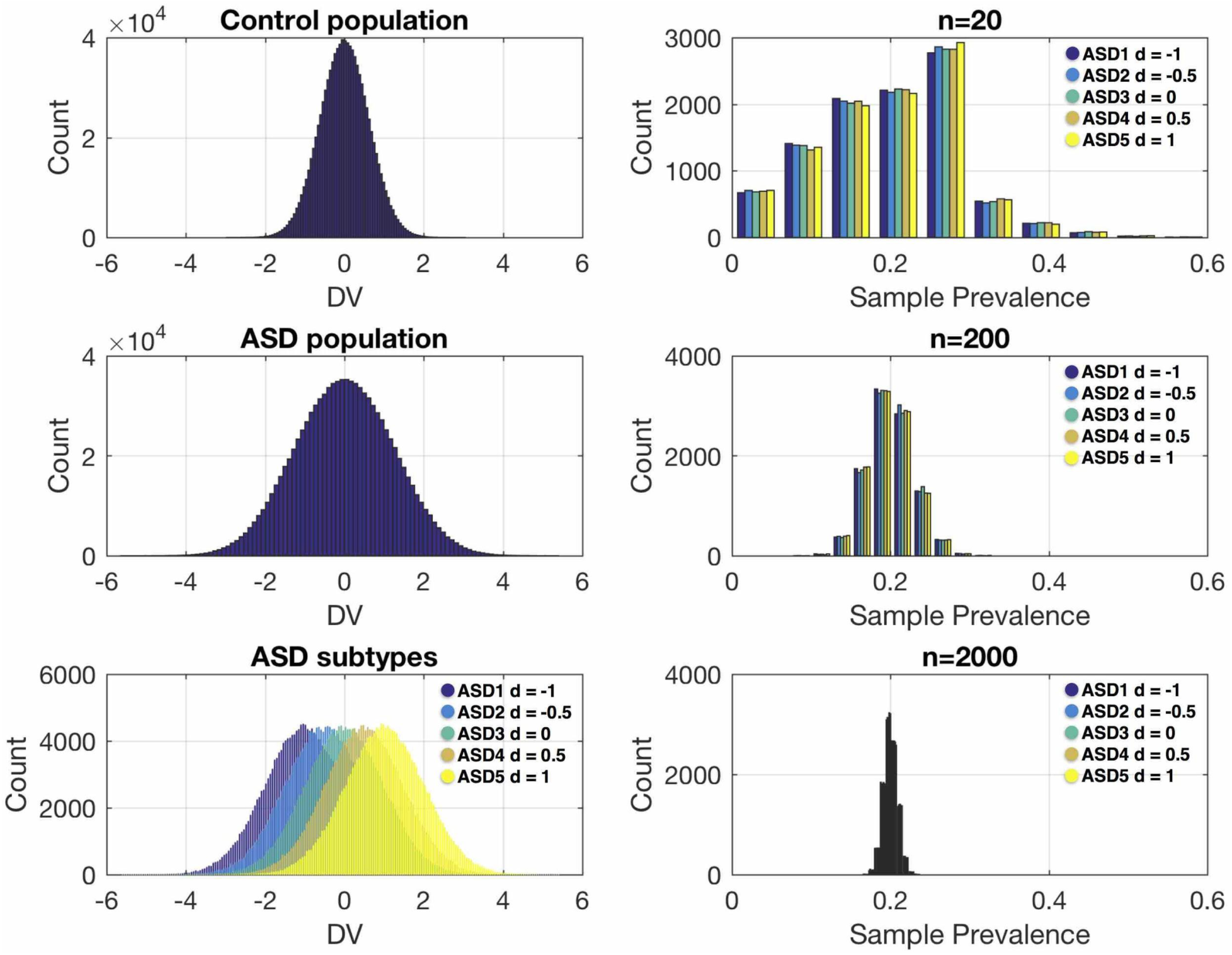
Simulation showing sampling variability and bias of enrichment of specific strata in small sample size studies. In this simulation we generated a control population (n=1,000,000) with a mean of 0 and a standard deviation of 1 on a hypothetical dependent variable (DV). We then generated an autism population (n=1,000,000) with 5 different autism subtypes each with a prevalence of 20% (e.g., n=200,000 for each subtype). These subtypes vary from the control population in effect size in units of 0.5 standard deviations, ranging from -1 to 1. This was done to simulate heterogeneity in the autism population that is reflective of very different types of effects. For example, the autism subtype 5 shows a pronounced increased response on the DV, whereas autism subtype 1 shows a pronounced decreased response on the DV. Across 10,000 simulated experiments, we then randomly sampled from the autism population sample sizes of n=20, n=200, and n=2000, and computed the sample prevalence of each autism subtype. The ideal result without any bias would be sample prevalences of around 20% for each subtype. This 20% sample prevalence is approached at n=2000, and to some extent at n=200. However, small sample sizes such as n=20 shows large variability in sample prevalences of the subtypes and this can markedly bias the results of a case-control comparison. The code for implementing and reproducing these simulations is available at https://github.com/mvlombardo/effectsizesim.

Such biases due to sampling variability across subtypes can paint a confusing picture and has considerable importance for replicability. To illustrate, we give a simple example indicative of many cases in the current literature. For example, Study 1 may unknowingly possesses a sample enriched with specific autism subtypes that show a decreased response on some dependent variable. Study 2 unknowingly has a different autism sample enriched with subtypes that show a contradictory increased response on the same dependent variable. Both studies are published and the authors of each may get into a heated debate, claiming that the other has the wrong viewpoint. Yet a third study comes out with perhaps a more unbiased (and possibly larger) sample, and given that the overall population effect could be near zero for a case-control comparison (as in the simulations in Fig 4), this third study finds no difference and claims that both studies 1 and 2 are false positives. While the third study may be the clearest indication of what occurs as an overall case-control effect, this study too may be missing the point completely – the population under investigation is not homogeneous and is stratified. Therefore, each study could have merit, if better contextualized and with some attempt to grapple with issues of heterogeneity. Thus, it is clear from these example that practices of running case-control studies, utilizing small sample sizes, and not fully confronting the issue of heterogeneity in autism may compound our problems and lead to a conflicted literature and the possibility of delayed scientific progress.

Given these considerations, our recommendation is to shift away from case-control models and more towards stratified and/or dimensional models that take into account important heterogeneity in autism. However, the suggestion here is easier said than done in practice. Thus, it may be important to spell out some reasons for why we think case-control models still persist. Conducting studies with very large sample sizes is challenging and perhaps only the most well-funded laboratories and/or consortiums can regularly conduct such work. In a situation where we are investigating phenomena with stratified models, this problem is magnified since one now needs large sample sizes within each strata being investigated. The practical issues are further compounded when there is need to replicate – a need which we would fully suggest is absolutely necessary to build confidence in identified effects. Therefore, it is easy to see why in practice the case-control paradigm persists, due to its practical ease compared to a stratified or dimensional model. It is also the case that many interesting types of stratifications may not be apparent to investigators and this could be another reason for the persistence of case-control models. Certainly there is also some value in results from well done work utilizing case-control models. Nevertheless, in order to make substantial progress towards precision medicine, we should begin shifting paradigms towards more research utilizing stratified and/or dimensional models that explicitly try to explain aspects of heterogeneity in autism. Contributions towards delineating heterogeneity could still be made by studies with moderate-sized samples, as long as a stratified model is applied in an *a priori* manner and with sufficient statistical power and ideally demonstration of replication. Such studies could focus on examining narrowly defined subgroups in the autism spectrum, derived either from hypothesis-driven strata (e.g. individuals with specific behavioral profile, specific neurobiological status, specific developmental characteristics, specific etiological factor, etc.) or strata discovered via prior data-driven studies.

## Essential big data characteristics for studying heterogeneity

While the idea of heterogeneity in autism has been around for some time, it is understandable why as a field we have made only limited progress. Conducting research on heterogeneity can be difficult for reasons of lack of data. As the previous discussions on issues with small sample sizes suggest, we would argue that one key ingredient to successfully studying heterogeneity in autism is ‘*big data*’. When we use the phrase ‘*big data*’, we are not necessarily referring to the ‘feature’ dimension of the data – that is, massively multivariate ‘feature rich’ data (e.g., neuroimaging or genomics data). Obviously, feature rich aspects of big data are indeed important in their own right and for the purposes of understanding heterogeneity. Rather, the dimensions we would emphasize about big data are the participant dimension (i.e. large sample size) as well as the depth of the measured features embedded in the participant dimension. Put another way, we need big data that has characteristics of being is both ‘*broad*’ and ‘*deep*’^75^ (Fig 1C).

*Broad* data refers directly to participant or sample dimension of the dataset (as opposed to the feature dimension) and is characteristic of massive sample size. Such a broad spread over individuals should ideally provide good coverage over the population of interest and allows for sufficient sampling of each strata of interest. Broad data is, we argue, an essential big data ingredient for decomposing heterogeneity in autism since, as noted above, we can run into many problems with data that is not sufficiently large or does not allow for such broad coverage over the population. Sufficiently broad data can also open up opportunities for replicating findings, since experimental designs can be planned out to hold ahead of time to set aside a sufficiently large validation set to replicate findings from an initial broad discovery set. As data sharing and open data initiatives become more prevalent, we should see more investigations on heterogeneity that meet this big data requirement. There are some current resources that are immediately available to meet such needs (e.g., the ABIDE datasets^76^, the National Database for Autism Research (NDAR)^77^, the Simons Simplex Collection^78^, SPARK^79^, the Healthy Brain Network^80^, and see^81, 82^) and we would expect much more in the coming years. As we get better at detecting what are the relevant dimensions and/or subtypes explaining important heterogeneity in autism, we may be better able to design high-powered targeted studies where the requirements for massive n may be reduced substantially. However, for most topics, we are not yet at this stage, and thus, broad data with massive sample size is necessary.

Developing models to explain aspects of heterogeneity at one level is only the first step. Once we have built good models that explain heterogeneity at one level, we will need to ask the next translational question: ‘*What else are these models good for?*’ Put differently, stratified or dimensional models can be good at predicting phenomena at one level of analysis, but because autism is heterogeneous at multiple levels, could such models help us make sense of heterogeneity outside of the domain that the model was originally built upon? Answering this question can have considerable impact towards precision medicine goals. For instance, a geneticist may have identified a unique biological subtype of autism based around a certain genetic mechanism. Such a genetic stratifier would already be quite useful for pinpointing a specific discrete cause for some proportion of the autism population. However, working towards precision medicine, we would next want to know whether such a genetic subtype is different from other autistic individuals on clinically relevant aspects such as prognosis, response to treatment, symptomatology, cognition, etc. Thus, when we ask this type of question, we need big data that is not only broad, but also ‘*deep*’^75^. Deep data is data collected on the same individuals that penetrates through multiple levels of analysis (Fig 1C). Deep data allows for stratifications or dimensional models to be built at one level, but the important tests of such stratifications can be done at other levels. An example of this can be seen in recent work on the Simons Simplex Collection. Here the authors made stratifications on the phenotype and then asked the question of whether such stratifications increased power for detecting GWAS-type effects at the genetic level^30^. Thus, to best answer questions by utilizing stratified or dimensional models, we will require big data that is both broad and deep, as the combination of both types of data can allow for discovery of explanations of autism heterogeneity and can immediately point towards the utility of such models for explaining the multi-level complexity inherent in autism. New multi-site studies such as EU-AIMS LEAP are targeted to directly address both issues of broad and deep data^83-85^ and we need other efforts along these lines.

## Approaches to decomposing heterogeneity in autism: top-down, bottom-up, and chronogeneity

Since the approach to decomposing heterogeneity in autism towards precision medicine goals is one of identifying clinically and mechanistically useful models, it is helpful to make salient some different approaches towards these goals. A common circumstance might be where a researcher makes a stratification at a level higher up in the hierarchy presented in Fig 1C. The translational next step may be to work down towards understanding how a stratified and/or dimensional model at this higher level of analysis can explain some phenomenon at a lower level. We refer to this as a *top-down* approach. For example, a clinically-important stratification can be made in the early development of autism regarding language outcome at 4-5 years of age. Some children keep up with age-appropriate norms in the areas of expressive and receptive language development, whereas others fall far behind in their language abilities across these domains. The empirical question after making such stratification could be whether such autism language-outcome subtypes differentiate at the level of neural systems organization, particularly neural systems that are developing specialization of function for speech and language processes^22^. In this example, it is clear that the stratifications were made at a level of analysis above the level that was later interrogated for mechanistic utility. Thus, while on its own, early language outcome was itself a clinically important stratifier, this top-down work also indicates that the stratifier may also be mechanistically useful for pointing towards different underlying biology. Other examples of a top-down approach may be based on cognitive characteristics^86^, sex/gender^74^, and co-occurring medical and psychiatric conditions (e.g. epilepsy^87^, ADHD^26^, etc). This type of top-down approach may then ultimately motivate future work that could potentially identify unique discoveries about biology behind a subset of the autism population that was previously unknown.

In contrast to top-down approaches, an approach that works from the *bottom-up* could be highly complementary. As the phrase implies, a *bottom-up* approach starts with identifying and building useful models from a lower level in the hierarchy, and then asks questions about how such low-level models can explain phenomena higher up in the hierarchy. For example, in the ‘*genetics first*’ approach, an investigator may be interested in identifying how different high-impact genetic causes for autism may be similar or different at a phenotypic or cognitive level of analysis^88-91^. In another example, an investigator may compare autism subtypes at the level of neural systems or structural brain features (e.g., with or without early brain enlargement), and then ask the question of whether such a stratification provides a meaningful indicator of differentiation at a clinical level^14^. Both top-down and bottom-up approaches can be useful, depending on the particular research question, and each can highlight different aspects of important heterogeneity in autism. In order to link up such multi-level complexity into explanations behind heterogeneity in autism, it will be imperative for work to engage in both of these approaches.

A final approach to decomposing heterogeneity deals with the lifespan developmental dimension across any level of analysis, or ‘chronogeneity’^37^. Several large longitudinal studies consistently indicate that there are several autism subtypes with different developmental trajectories^16-18, 37, 92^. Regression, a developmental feature not uncommonly seen in autistic individuals, is another key stratifier that is surprisingly under-studied but with plausible unique biological bases^93, 94^. Within the developmental dimension, heterogeneity can be assessed as both inter-and intra-individual variability, but can also cover individualized deviance from group trajectories over time^37^ or age-specific norms^95, 96^. Chronogeneity thus offers a unique vantage point on multi-level heterogeneity not covered by understanding heterogeneity at static time points.

## Approaches to decomposing heterogeneity in autism: supervised versus unsupervised

In addition to conceptualizing stratified and/or dimensional models by top-down, bottom-up, or developmental approaches, it is also important to clarify how we build on the process of understanding heterogeneity. Ultimately, the scientific process of better understanding heterogeneity in autism is a learning problem. Taking ideas from statistical or machine learning, we can broadly divide learning processes into *supervised* and *unsupervised* learning^97^. Supervised learning deals with *a priori* knowledge about a topic (i.e. known labels), and then seeks to derive a model to best predict that known information. With regard to the process of understanding heterogeneity in autism, the analogy of supervised learning can apply to all instances where the experimenter uses their own knowledge and justifications to dictate where the stratifications are made (e.g., top-down, bottom-up, or developmental). In other words, knowledge from a supervised source (e.g., an investigator, a theory) informs the stratification or dimension to be modeled. This type of approach has the advantage of being theory-driven and/or builds on expert knowledge of the investigator (e.g., clinical intuition or experience), who may already have highlighted a distinction that is meaningful and justified in a variety of ways.

The disadvantage of solely relying on a *‘supervised’* approach is that the investigator and/or a theory may be missing other important distinctions about how to model heterogeneity for the question of interest. In this case, the learning process can be helped by some type of *‘unsupervised’* statistical learning process that uncovers distinctions that may not be readily apparent from *a priori* knowledge. Because big data is a key ingredient for building models to explain heterogeneity, we can utilize the feature-rich aspects of big data to embark on data-driven discovery of potentially complex multivariate patterns that distinguish different types of individuals. We refer to this data-driven approach as an *‘unsupervised’* approach since computationally, the learning occurs without any expert *a priori* knowledge and justifications and solely relies on statistical distinctions embedded in the data itself. With this approach we likely rely on advanced computational techniques from machine learning that are tailored to best identify complex multivariate distinctions. For example, we utilized clustering methods taken from systems biology and applied them to item-level patterning of behavioral responses on the Reading the Mind in the Eyes Test (RMET). This unsupervised approach yielded discovery of 5 different autism subtypes that could be replicably identified in an independent replication set (Fig 2)^24^. In other work, Ellegood and colleagues applied clustering to neuroanatomical phenotypes across a range of different mouse models for autism. This work illustrated that heterogeneous starting points (e.g., different genetic mutations highly associated with autism) can converge and diverge at the level of neuroanatomical phenotypes^98^. Feczko and colleagues utilized a community detection algorithm, typically used in network science to discover ‘modules’, to detect subgroups of autism based on a variety of measures from a neuropsychological test battery. This cognitive subtyping was then followed by examination and identification of different patterns of resting state functional connectivity in the brain that were driven primarily by subgroups^15^. Using structural MRI measures of cortical morphometry, Hong and colleagues used clustering to identify 3 autism subtypes with different anatomical profiles. These anatomically-defined subtypes were then found to be useful for increasing the performance of supervised learning models to predict symptom severity on measures such as the ADOS^12^.

It should be noted that both supervised and unsupervised approaches have their advantages and disadvantages, and can be complementary. An example of this complementarity can be seen in a hybrid supervised/unsupervised approach from Feczko and colleagues^15^. In this study, the authors utilized a supervised ensemble learning model called Functional Random Forest (FRF), to classify autism versus typically developing children based on cognitive features from a neuropsychological test battery. In addition to classifying autism versus typically developing children, the FRF model produces a proximity matrix that indicates similarity between individuals. The authors then utilized this proximity matrix to identify subgroups in an unsupervised manner utilizing a community detection algorithm, typically used in network science to discover ‘modules’. This hybrid approach to cognitive subtyping proved useful as a top-down approach towards identifying different patterns of resting state functional connectivity across the subtypes. Thus, through the scientific process of building knowledge about important stratified or dimensional models, both unsupervised and supervised approaches can inform each other, and in some cases may be utilized together in a hybrid fashion.

## Decomposing heterogeneity in relation to transdiagnostic constructs

Although so far we treat autism as an entity and focus on heterogeneity within it, this diagnostic construct is human-made, cumulative and evolving^99, 100^. Phenotypically, autism frequently co-occurs with other neurodevelopmental (e.g., ADHD, tic disorders) and psychiatric (e.g., anxiety, depression, obsessive-compulsive disorder, psychotic disorders) conditions^1, 101^ and heightened autistic traits often cut across other categorical diagnoses as well^102^. Underlying this may be multi-level processes cutting across sets of frequently co-occurring diagnoses^103^, which potentially can be delineated by transdiagnostic approaches such as using the RDoC framework^46^. In this respect, we should acknowledge that heterogeneity in autism is part of the broader heterogeneity existing across neurodevelopmental and psychiatric conditions. In the same vein, the reasons, principles and approaches described above to tackle heterogeneity in autism can be similarly applied when autism is studied within a transdiagnostic framework cutting across multiple diagnoses. In the background of high co-occurrence, a transdiagnostic framework is necessary in deepening our understanding of the heterogeneity within and beyond autism.

### Conclusions

Understanding how heterogeneity manifests in different individuals with autism is amongst the biggest challenge our field currently faces. As we continue to develop models for explaining this heterogeneity, it will be useful to capture such work under the organizing concepts we have laid out in this article. Heterogeneity must be interpreted relative to the zeitgeist, particularly as it pertains to how diagnostic concepts evolve. Models for explaining heterogeneity manifest in many ways, depending on whether the researcher conceptualizes the differences between individuals as quantitative and dimensional, or qualitative and categorical. There is room for both models that fuse together both dimensional and categorical distinctions. In general, we need to move beyond one-size-fits-all models such as case-control models, and we need to be stringent with respect to methodology, since practices such as small sample size research cannot live up to the challenges that heterogeneity creates. Small samples cannot adequately cover heterogeneity in the autism population in a highly generalizable fashion, and hence there is a need for ‘big data’ when studying heterogeneity. Big data should be both broad and deep, to sample adequately across different strata from the population but also to examine how strata defined at one level may be relevant for explaining variability at other levels. Heterogeneity can be parsed from multiple approaches that capitalize on information from levels of analysis either most proximate or most distal from the clinical phenotype and which work their way down or up through the hierarchy, or via examination of changes over development. Also important for conceptually organizing work on this topic is whether we utilize *a priori* knowledge to build heterogeneity models or whether we allow computational methods to inform us about data-driven distinctions that may be hidden and not readily apparent to most researchers. Finally, models to understand heterogeneity can move beyond just those with clinical diagnoses of autism and, in the future, transdiagnostic approaches utilizing similar organizing concepts may provide complimentary information. Overall, the push to understand heterogeneity is critical as we attempt to move towards precision medicine.

## Acknowledgments

MVL was supported by a European Research Council (ERC) Starting Grant (ERC-2017-STG 755816). M-CL was supported by the O’Brien Scholars Program within the Child and Youth Mental Health Collaborative at the Centre for Addiction and Mental Health (CAMH) and The Hospital for Sick Children, Toronto, and the Slaight Family Child and Youth Mental Health Innovation Fund, CAMH Foundation. SBC was supported by the Autism Research Trust and the Medical Research Council during the period of this work. The research was supported by the National Institute for Health Research (NIHR) Collaboration for Leadership in Applied Health Research and Care East of England at Cambridgeshire and Peterborough NHS Foundation Trust. The views expressed are those of the authors and not necessarily those of the NHS, the NIHR or the Department of Health.

## Disclosures

All authors have no conflict of interests to declare.

